# Early feature extraction drives model performance in high-resolution chromatin accessibility prediction: A systematic evaluation of deep learning architectures

**DOI:** 10.1101/2025.03.01.641000

**Authors:** Aayush Grover, Till Muser, Liine Kasak, Lin Zhang, Ekaterina Krymova, Valentina Boeva

## Abstract

Fine-grained prediction of chromatin accessibility from DNA sequence is a foundational step in modeling gene expression changes resulting from sequence variants. Yet, few methods operate at the resolution necessary to capture subtle effects of single-nucleotide changes. Furthermore, it remains unclear which architectural components—such as residual connections, normalization strategies, or attention mechanisms—drive performance in these high-resolution predictions. To address these knowledge gaps, we systematically evaluate classic architectural choices and introduce ConvNeXt V2 blocks, originally developed for computer vision, as high-resolution feature extractors in deep learning models for genomic data. Integrated into diverse architectures—CNNs, LSTMs, dilated CNNs, and transformers—ConvNeXt V2 blocks consistently improve performance, leading to similar prediction accuracy across these different model types. This reveals that early feature extraction, rather than downstream architecture, is the primary determinant of prediction accuracy. A comprehensive evaluation of these models on ATAC-seq signal prediction at 4 bp resolution in a cell type-specific manner identifies the ConvNeXt-based dilated CNN as the most robust performer, better preserving the signal’s shape. Our codebase and benchmarks provide practical tools for high-resolution chromatin modeling.

## 1 Introduction

Predicting chromatin accessibility from DNA sequence is one of the central challenges in regulatory genomics, with applications including the interpretation of non-coding genetic variants and the decoding of regulatory element architecture [1–4]. Accurate, high-resolution maps of chromatin accessibility are essential for downstream tasks such as identifying regulatory elements, modeling 3D chromatin organization, and predicting gene expression [5–9]. While experimental assays like FAIRE-seq [10], DNase-seq [11], and ATAC-seq [12] provide high-resolution measurements of chromatin accessibility in specific cellular contexts, they are resource-intensive and often unavailable for specific genetic variants. In the absence of such experimental assays, neural network–based predictions offer a reliable and efficient alternative for estimating changes in chromatin accessibility induced by the genetic variants in a cell type-specific manner.

Numerous deep learning approaches have been proposed to predict chromatin accessibility from DNA sequence. Earlier methods [1, 13–16] treated this as a binary classification task, labeling regions as either open or closed. Some of the later methods, such as Deopen [17] demonstrated the performance of their model in predicting the maximum intensity of chromatin accessibility corresponding to the input DNA segment. Subsequently, Basenji [2] and Enformer [3] showed that it was possible to predict the intensity of the chromatin accessibility signal corresponding to the input DNA sequence with a relatively lower resolution (128bp) by binning the signal tracks.

Although these methods are highly accurate, this coarse formulation (*i*.*e*., predicting chromatin accessibility profiles at 128bp or lower resolution) is still insufficient to study local changes in chromatin accessibility caused by point mutations. Moreover, it hinders their applicability for downstream tasks, including prediction of transcription factor binding [18], cell-type-specific chromatin structure [6, 7, 9, 19, 20], and gene expression [8, 9]. Therefore, there is a need for the development of deep learning methods that can accurately predict high-resolution chromatin accessibility signals.

Recently introduced ChromBPNet [4] predicts chromatin accessibility at base-pair resolution using dilated convolutional networks, representing an advance in resolution capabilities. However, it is not extensively compared against other model architectures, like recurrent neural networks and transformers, leaving questions about optimal architectural choices unanswered. GOPHER [21] was introduced as an extensive benchmark that compared various deep learning methods, such as Basenji [2] and BPNet [22], in their ability to predict chromatin accessibility for different experimental setups. Both of these methods use dilated convolutional neural networks (dCNNs) to capture long-range information from the input DNA sequence. These methods were evaluated against a convolutional neural network (CNN) baseline, highlighting the predictive prowess of dCNN-based methods over CNN-based methods. However, recurrent neural network-based methods and transformer-based methods are not explored in this benchmark, creating a gap in understanding which architectural paradigms perform best for high-resolution chromatin accessibility prediction.

In this work, we present a systematic evaluation of deep learning models for predicting ATAC-seq signals at 4 bp resolution from DNA sequence. We begin by extensively tuning three commonly used architectures—CNNs, recurrent neural networks (specifically, long short-term memory, LSTM), and dCNNs—for high-resolution chromatin accessibility prediction. Building on this, we incorporate ConvNeXt V2 [23] blocks, originally developed for vision tasks, as feature extractors across all architectures and introduce a novel transformer-based model tailored for this task. We evaluate each model’s performance on accessible regions and genome-wide, assess robustness to input shifts, and test sensitivity to single-nucleotide variants (SNVs) using cancer patient data. This benchmark reveals how architectural choices impact predictive accuracy, robustness, and variant interpretation, providing a practical foundation for future development of deep learning methods in regulatory genomics. The documented code of our ASAP framework for Allele-Specific ATAC-seq Prediction is available at https://github.com/BoevaLab/ASAP.

## 2 Results

### 2.1 Designing a deep-learning model benchmark for high-resolution ATAC-seq prediction

We evaluated four categories of deep learning architectures—convolutional neural networks (CNNs), recurrent neural networks (LSTMs), dilated CNNs (dCNNs), and transformers—based on their ability to predict ATAC-seq signal at high resolution. To ensure a fair comparison, we standardized the experimental setup across models: each received a 2048 bp one-hot encoded DNA sequence as input and was trained to predict the chromatin accessibility signal at 4 bp resolution across the central 1024 bp region (Fig. 1). This resolution was chosen because transcription factor binding motifs typically span more than 4 bp [24], making it sufficient to capture relevant regulatory variation. The 2048 bp input window also provides adequate genomic context for the models to learn both local and distal sequence features.

**Fig. 1.**
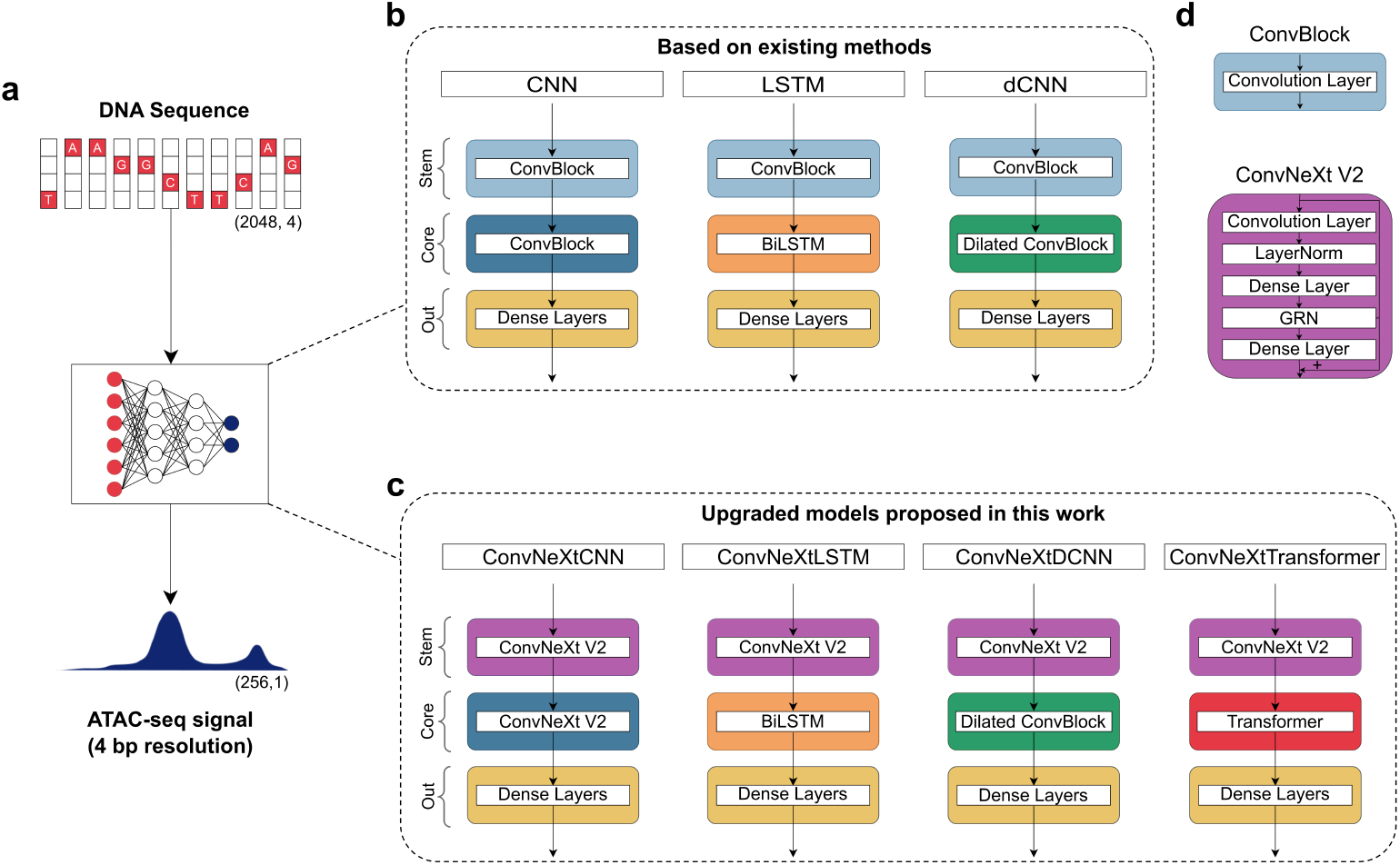
Overview of the benchmarking setup. **a**, Models are compared based on their ability to predict experimental ATAC-seq at 4 bp resolution from an input of 2048 bp long DNA sequences. **b**, Models based on existing methods include CNN, LSTM, and dCNN-based architectures. Each model has a convolution-based stem to extract genomic features. **c**, Models proposed in this work, including a transformer-based architecture. The new models use ConvNeXt V2 blocks to effectively extract features from the input DNA sequence. **d**, ConvBlock uses a single convolutional layer whereas ConvNeXt V2 block additionally uses dense layers, layer normalization and Global Response Normalization (GRN).

We began by benchmarking architectures used in existing methods, broadly categorized into CNNs [1], LSTMs [15, 17], and dilated CNNs [2, 4] (Fig. 1a). Since these models were originally developed for different prediction tasks and resolutions, we conducted an extensive hyperparameter search to adapt each to our standardized 4 bp resolution setup. Next, to improve feature extraction, we replaced standard convolutional layers in the stem of all three architectures with transformer-inspired ConvNeXt V2 blocks, originally designed for vision tasks but well-suited for capturing local and global genomic features [23] (Fig. 1b).

Previous studies have shown that transformer architectures can outperform convolution-based models in capturing long-range dependencies in genomic sequences [3, 25–27]. However, their application to high-resolution chromatin accessibility prediction has not been explored. To address this limitation, we designed a novel transformer-based model incorporating rotary positional embeddings [28] (Fig. 1b).

The detailed architecture of each model can be found in Supplementary Fig. S1. Together, these models form a comprehensive benchmark across four deep learning architecture families.

### 2.2 Improving model prediction accuracy with ConvNeXt-based feature extraction

Each model was extensively trained and tested on data from eight distinct datasets: four established human cell lines (GM12878, K562, IMR90, and HepG2) and four cancer patient samples (two colon adenocarcinomas (COAD) and two kidney renal papillary cell carcinomas (KIRP)). We implemented a cell type-specific training approach to account for the distinct regulatory mechanisms that govern chromatin accessibility in different cellular contexts. To ensure statistical robustness, we implemented a 5-fold cross-validation strategy for each model within each cell type or tumor sample. The validation folds were stratified by chromosomes, allowing us to evaluate model performance on entirely unseen genomic regions while maintaining the unique characteristics of each cell type.

Within the test chromosomes, we evaluated model performance using two complementary measurements: prediction accuracy on the reported ATAC-seq peaks and prediction accuracy on entire genomic regions. This dual assessment allowed us to benchmark both the models’ ability to accurately characterize highly accessible chromatin regions (peak-focused evaluation) and their capacity to distinguish between accessible and inaccessible regions across the complete chromosomal landscape (genome-wide evaluation).

Among the architectures employed in existing methods (CNNs, LSTMs, dCNNs), dCNNs consistently outperformed others in both peak-focused and genome-wide settings (Fig. 2a, Supplementary Fig. S2). However, upon incorporating ConvNeXt V2 blocks in the stem of these architectures, all models demonstrated remarkably similar high correlations with the experimental ATAC-seq values in the accessible regions as well as on whole genome (average Pearson’s R *≥* 0.7) (Fig. 2b, Supplementary Fig. S2).

**Fig. 2.**
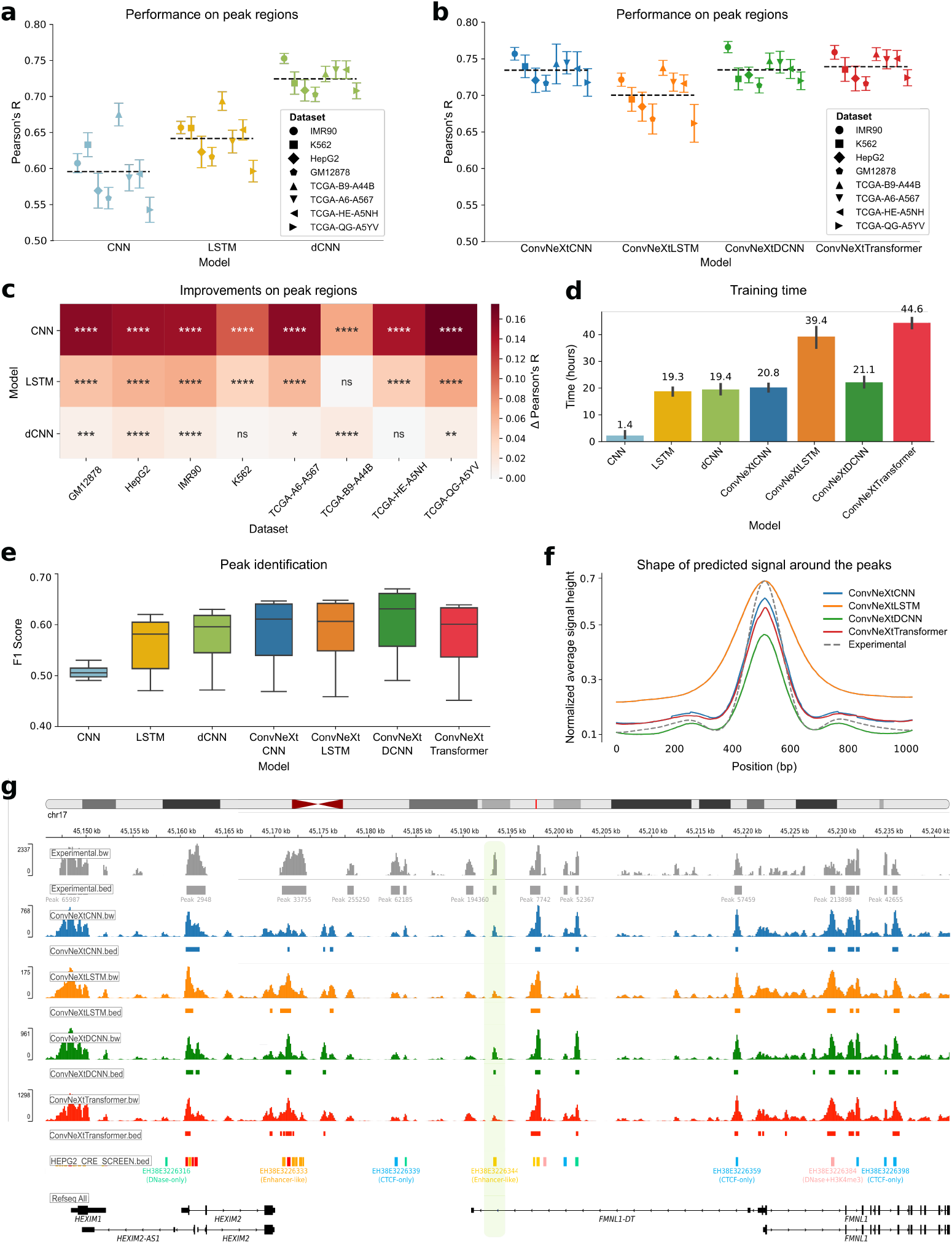
Model comparison across datasets. **a**, Pearson’s correlation between true and predicted ATAC-seq signals in peak regions across eight distinct datasets for state-of-the-art models. **b**, Performance comparison among the four models proposed in this work (ConvNeXtCNNs, ConvNeXtLSTMs, ConvNeXtDCNNs, and ConvNeXtTransformers) for the ATAC-seq peak regions stratified by cell lines and primary tumor samples. Each point depicts a test chromosome. The black dashed line shows the average performance of a model across all datasets and chromosomes. **c**, Improvements of the new ConvNeXt-based methods proposed in this work for the ATAC-seq peaks as compared to existing methods. The significance is calculated with a two-sided Mann-Whitney U test on Pearson’s R calculated for each test chromosome. ****: *P ≤* 0.0001, ***: *P ≤* 0.001, **: *P ≤* 0.01, *: *P ≤* 0.05, ns: *P >* 0.05. The Δ Pearson’s R is calculated as the difference between mean Pearson’s R across all chromosomes for a ConvNeXt-based method and the corresponding existing method. **d**, The total training time (in hours) of each method on a single RTX2080Ti GPU averaged across eight datasets. The error bar shows 95% confidence interval. Median training time is mentioned over each bar. **e**, The F1 score is calculated for all the methods to evaluate the retrieval of ATAC-seq peak calls from the predictions. **f**, The average shape of the predicted ATAC-seq signal is compared against the experimental shape in the ATAC-seq peak regions. **g**, An example of predicted signal vs. the experimental signal around the *FMNL1* gene in HepG2 cells. A distal enhancer, discovered as a peak only based on the predictions of the ConvNeXtDCNN method, is highlighted.

The novel transformer-based architecture, ConvNeXtTransformer, that we also introduced in this evaluation, performed comparably to the ConvNeXtCNN and ConvNeXtDCNN models. We also explored the xLSTM architecture, a more recent development than the transformer, which offers improved memory efficiency for sequential data processing [29]. However, when tested in our framework, ConvNeXt-xLSTM was outperformed by ConvNeXtLSTM, suggesting that its recurrent, long-range dependency modeling impaired the capture of local context critical for ATAC-seq prediction compared to ConvNeXtTransformer’s attention-based approach (Supplementary Fig. S3). Overall, the addition of the ConvNeXt blocks to the stem significantly improved the performance of the existing models across all datasets, highlighting the better effectiveness of ConvNeXt blocks in capturing genomic features with local and global context (Fig. 2c, Supplementary Fig. S2).

Adding ConvNeXt blocks to the stem, however, increased training times, with ConvNeXtDCNN requiring approximately 1.7 more hours as compared to dCNN (Fig. 2d). Despite this, all models exhibited fast inference, predicting ATAC-seq signal across the entire chromosome 17 in under 2 minutes, underscoring their practical utility for variant-effect prediction tasks (Supplementary Fig. S2).

### 2.3 Assessing peak detection and signal shape fidelity in predicted ATAC-seq tracks

Accurate identification of ATAC-seq peaks is essential for deep learning models to generate outputs suitable for downstream tasks. We performed peak calling on the predicted ATAC-seq signals from each model (Methods) and compared the resulting peaks against experimental data. For robust evaluation, we used the F1 score, which accounts for class imbalance. ConvNeXt-based models outperformed existing baselines, with ConvNeXtDCNN achieving the highest F1 score (Fig. 2e).

To assess how well these models capture signal shapes around the peak maxima, we plotted the min-max normalized predicted ATAC-seq signal averaged across all peaks, datasets, and training folds (Fig. 2f). While ConvNeXtLSTM tended to over-predict the signal at peak boundaries, producing smoother profiles, other ConvNeXt-based methods closely matched the experimental ATAC-seq signal, accurately capturing nucleosome positioning around peak centers.

Finally, we visualized the chromatin accessibility signals generated by each ConvNeXt-based model on test chromosomes and their corresponding peaks (Fig. 2g). The predicted signals and peaks co-localized with many known promoters and enhancers [30], corroborating quantitative overlap analyses with ground-truth accessible regions and confirming the high fidelity of predicted peak shapes (Fig. 2e-f). Notably, some enhancers were uniquely identified by specific methods; for example, an enhancer in the gene body of *FMNL1-DT* in HepG2 cells was marked as accessible only by ConvNeXtDCNN (Fig. 2g). Overall, ConvNeXtDCNN was the top-performing method based on its peak-calling accuracy and correlation with experimental data, with other ConvNeXt-based models also showing strong performance.

### 2.4 Evaluating model robustness to shifts in the DNA sequence inputs

A critical property of reliable genomic models is robustness to input shifts, *i*.*e*., the ability to generate consistent predictions regardless of whether the input sequence is precisely centered on a peak maximum or slightly offset. To quantify this property, we performed a systematic robustness test on all ConvNeXt-based models, measuring the variation in predicted ATAC-seq signal across common genomic regions when the input window is shifted [21] (Supplementary Fig. S4). For each ATAC-seq peak across all test chromosomes and training folds, we generated 17 different input shifts and calculated the coefficient of variation across each 4 bp bin in the commonly predicted region. Lower coefficient of variation corresponded to higher robustness, indicating more consistent predictions despite slight shifts in input positioning.

The position-stratified coefficient of variation, averaged across all peaks in different datasets, showed that all methods displayed higher robustness at peak centers (Fig. 3a). The periodic variation pattern observed in ConvNeXtCNN can be attributed to the inherent sensitivity of convolutional networks to random input shifts [31].

**Fig. 3.**
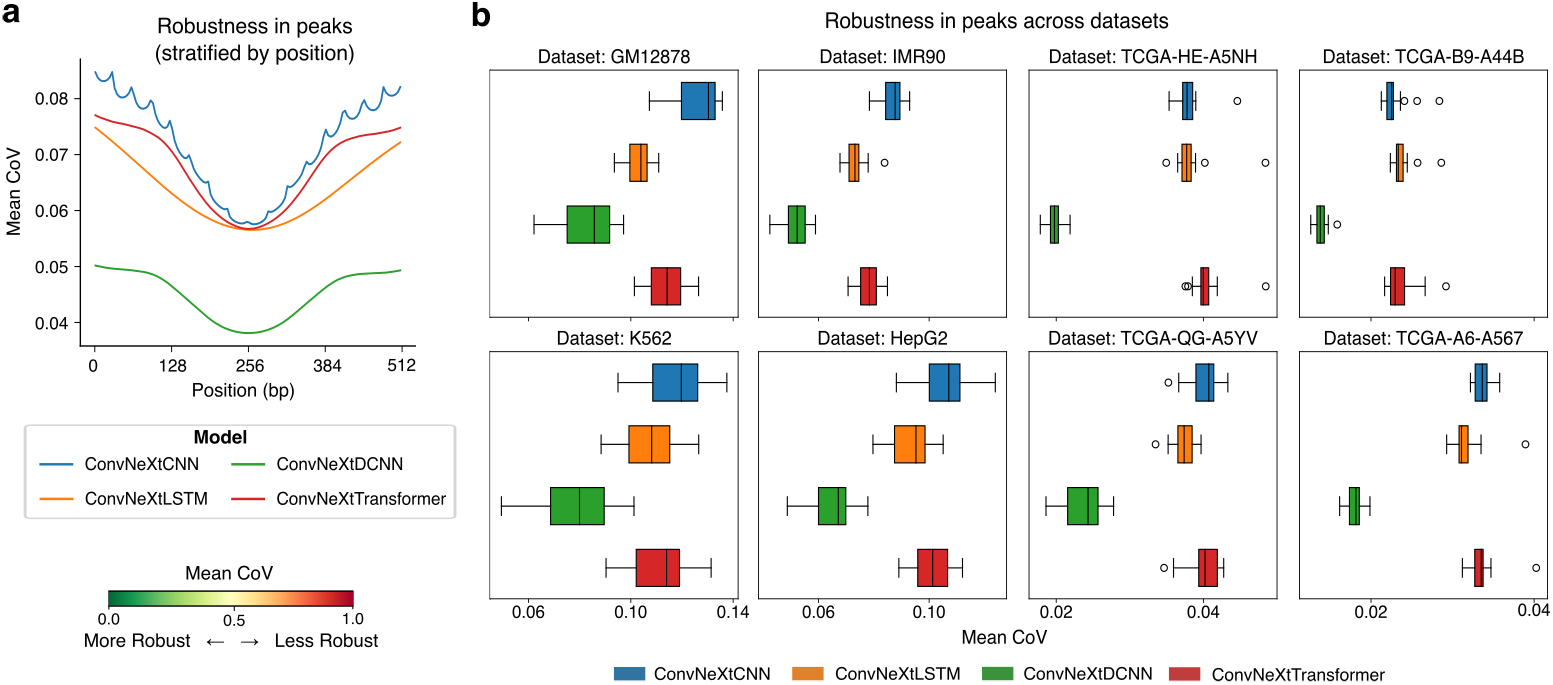
Evaluation of model robustness to input shifts. **a**, Position-stratified coefficient of variation (CoV) is calculated for all the methods for the ATAC-seq peak regions. The variation is averaged across all the peaks in unseen chromosomes of the 8 datasets. **b**, The mean CoV measured for ATAC-seq peak regions across the eight datasets used in this study. Each point represents mean CoV for a chromosome unseen by the model during training.

Among all the methods, ConvNeXtDCNN consistently emerged as the most robust method across all datasets, followed by ConvNeXtLSTM (Fig. 3b-c). The robustness of ConvNeXtDCNN was further confirmed by a low variation score observed using the entire unseen chromosomes of each dataset (Supplementary Fig. S4).

### 2.5 Evaluating model sensitivity to genetic point variants

Experimentally profiling chromatin accessibility for every genetic variant is prohibitively resource-intensive, highlighting the need for accurate in silico prediction of variant effects. To evaluate the ability of benchmarked models to capture the effects of genomic variants, we obtained SNVs from four cancer patients—two with colon adenocarcinoma and two with kidney renal papillary cell carcinoma—using data from the ICGC [32], and estimated variant-induced changes on chromatin accessibility based on ATAC-seq read counts (Fig. 4a). A total of 1,025 SNVs were analyzed after filtering out variants with minimal effect, defined as those with odds ratios between 0.67 and 1.33. For this task, all models were retrained on the full genome, excluding only regions containing SNVs. Model predictions were then generated using 2,048 bp SNV-centered input sequences, where the reference and variant sequences differed by a single nucleotide.

**Fig. 4.**
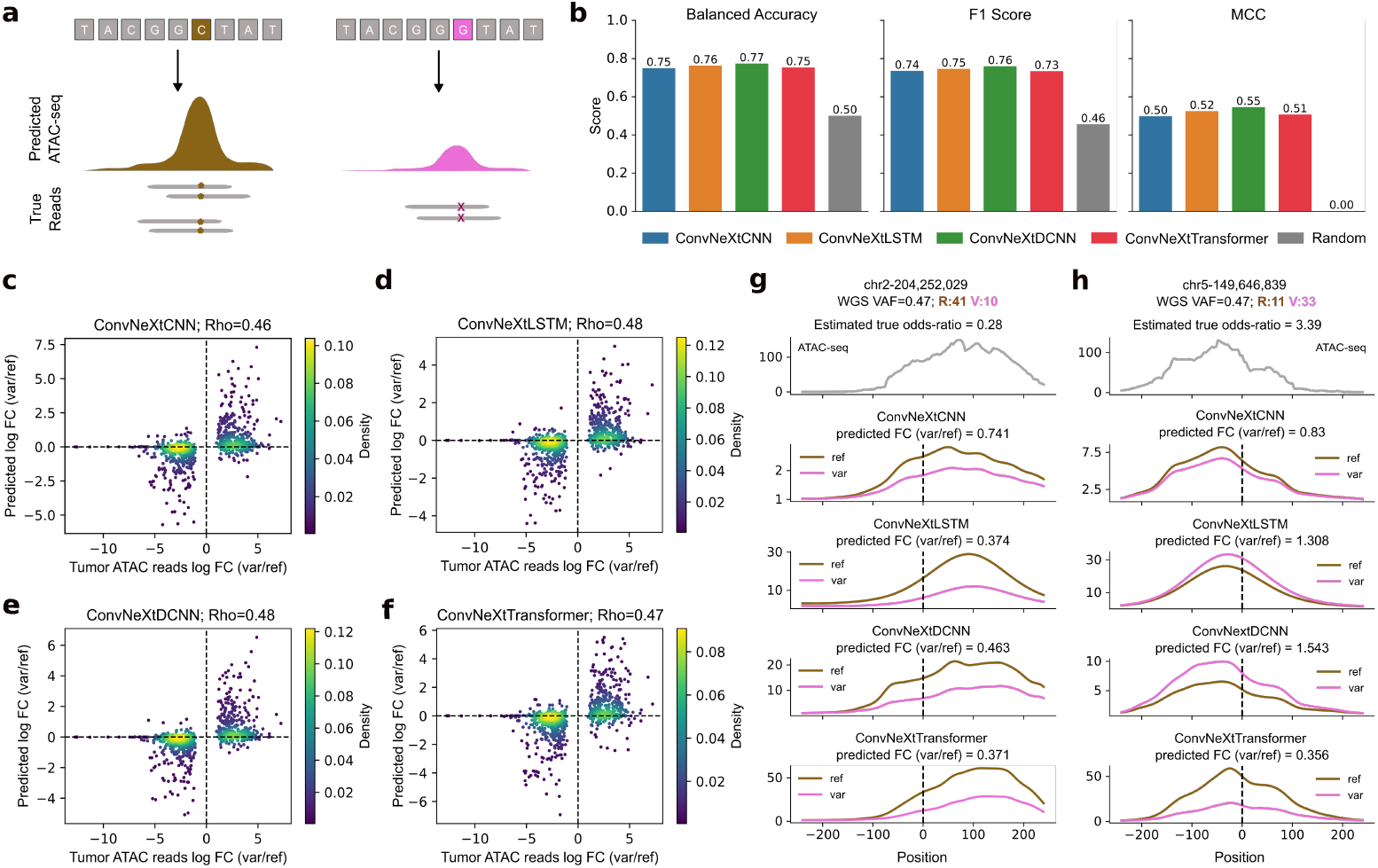
Evaluating model ability to accurately predict effects of single-nucleotide variants on chromatin accessibility. **a**, Experimental setup to test each model’s ability to predict the change in chromatin accessibility between the reference DNA sequence (left) and a genomic variant (right). **b**, ConvNeXt-based models are compared based on their ability to predict the correct directionality of change in accessibility between reference and variant. A total of 1,025 variants are considered, across all cancer patients included in this study. Balanced accuracy, F1 score, and Matthews correlation coefficient (MCC) are used as metrics. Random baseline is included to show the improvements in metrics by all the models proposed in this work. **c-f**, Scatter plots showing the quantitative change in accessibility by each model. Log-odds-ratio based on ATAC-seq read counts for reference and variants are compared against log-fold change (FC) between model predictions with reference and variant as model inputs and Spearman’s correlation (*Rho*) is calculated. **g-h**, Examples of allele-specific chromatin accessibility predictions for the reference allele and a genomic variant in chromosomes 2 and 5, respectively. WGS VAF: Whole genome sequencing variant allele frequency; R, V: ATAC-seq total reference reads and variant reads, respectively.

We first assessed the ability of each model to correctly predict the directionality of change in chromatin accessibility between the reference and variant alleles. All models outperformed the random baseline. Moreover, ConvNeXtDCNN achieved the highest performance with balanced accuracy of 0.77, F1 score of 0.76, and Matthews correlation coefficient (MCC) of 0.55 (Fig. 4b). Both F1 score and MCC account for class imbalance and are, therefore, more robust metrics for this task.

We next tackled the more challenging task of quantitatively predicting the change in chromatin accessibility induced by an SNV. For this evaluation, the estimated logodds ratio from ATAC-seq read counts for the reference and variant alleles served as the ground truth. This was compared against the log-fold-change between the ATAC-seq signal predicted by each model for the variant and the reference nucleotide (Methods). ConvNeXtLSTM and ConvNeXtDCNN achieved the highest Spearman’s correlation with the true odds ratio (*Rho* = 0.48) (Fig. 4c-f), closely followed by ConvNeXtTransformer (*Rho* = 0.47) and ConvNeXtCNN (*Rho* = 0.46).

Although all models mostly agreed on the predicted directionality of change in ATAC-seq signal, for many SNVs, ConvNeXtDCNN most accurately captured the detailed shape of the predicted signal (Fig. 4g, Supplementary Fig. S5). An exception is however illustrated by an SNV near 149.6 Mb on chromosome 5, where only the ConvNeXtLSTM and ConvNeXtDCNN models correctly predicted the direction of change, with ConvNeXtDCNN yielding a prediction closer to the ground-truth odds ratio (Fig. 4h). In conclusion, while all models performed well on the task of predicting the effect of SNVs on chromatin accessibility, ConvNeXtDCNN had a slight edge over the other methods.

## 3 Discussion

In this work, we compared existing deep learning architectures—CNNs, recurrent neural networks (specifically, LSTMs), and dCNNs—on their ability to predict highresolution chromatin accessibility signals from genome sequences (Fig. 1a). To improve model performance, we proposed new ConvNeXt-based models: ConvNeXtCNN, ConvNeXtLSTM, and ConvNeXtDCNN. Additionally, we introduced a transformer-based model, ConvNeXtTransformer, to create a more comprehensive set of architectures for this task (Fig. 1b).

The ConvNeXt-based models outperformed their traditional counterparts in accurately predicting ATAC-seq signals at 4 bp resolution across eight different cell types and primary tumor samples. Among them, ConvNeXtCNN, ConvNeXtDCNN, and ConvNeXtTransformer most accurately captured the shape of the ATAC-seq peaks (Fig. 2). We further evaluated the robustness of these models to shifts in the DNA sequence inputs. ConvNeXtDCNN emerged as the most robust method, efficiently integrating local and global sequence features through its dilated CNN layers (Fig. 3). Finally, we assessed the model ability to predict changes in chromatin accessibility caused by single-nucleotide genomic variants. ConvNeXtLSTM and ConvNeXtDCNN achieved the highest accuracy in predicting the log-odds-ratio of ATAC-seq reads between variant and reference alleles. ConvNeXtDCNN also best captured the directionality of accessibility changes, accurately distinguishing between increases and decreases in chromatin accessibility upon SNV introductions (Fig. 4). This highlights the sensitivity of the ConvNeXtDCNN model to genomic variants and suggests its potential utility in studying the functional effects of non-coding genetic variants.

While our ConvNeXt-based models demonstrated strong performance in predicting chromatin accessibility and the effects of SNVs, our models were trained in a cell-typespecific manner. Therefore, they were optimized for individual cellular contexts rather than generalized across different cell types. While this design ensures high accuracy within a given cell type, it also means that a new model must be trained for each additional cell type of interest. Future work could explore data-efficient fine-tuning approaches to accelerate model adaptation to new cellular contexts without requiring full retraining.

Overall, our study highlighted the advantages of ConvNeXt-based models in predicting high-resolution chromatin accessibility and regulatory effects of genetic variants. By combining ConvNeXt blocks as genomic feature extractors with CNNs, LSTMs, dilated CNNs, and transformers, we established a comprehensive framework of sequence-based chromatin modeling. Our results demonstrated that ConvNeXtDCNN is not only highly accurate but also robust to input shifts and effective at capturing effects of SNVs on chromatin accessibility, making ConvNeXtDCNN a promising method for genome-wide regulatory analysis.

## Methods

### ATAC-seq bigwig data

We downloaded the ATAC-seq signal p-value tracks for four cell lines–GM12878, IMR90, HepG2, and K562, from the ENCODE Consortium [33]–where the experimentally measured ATAC-seq has been preprocessed with the open source processing pipeline [34]. These cell lines consist of healthy and cancer cells, thereby creating a diverse dataset.

To further diversify our validation datasets, we used ATAC-seq data of two colon adenocarcinoma (COAD) and two kidney renal papillary cell carcinoma (KIRP) tumors from The Cancer Genome Atlas (TCGA) [35]. These datasets have been made available by the Genome Data Commons (GDC) Data Portal [36]. The choice of these patients was made due to the availability of the data on their somatic mutations and single-nucleotide polymorphisms (SNPs) in the International Cancer Genome Consortium (ICGC) and GDC data portals, respectively [36, 37]. Moreover, these patients had the high number of combined SNV calls (*>* 175, 000) before applying SNV-filtering (described below) and high tumor purity (Consensus Purity Estimate *>* 0.8) [38].

The ATAC-seq BAM files for the primary tumor samples were downloaded from National Cancer Institute’s Genomic Data Commons portal [39]. Signal p-value BigWig files were then generated using the ENCODE’s ATAC-seq processing pipeline [34]. All datasets used in this study are listed in Supplementary Table S1.

### Preprocessing

For the ATAC-seq of each cell line, we composed a dataset for each cell line or cancer patient where each item is a pair of one-hot encoded DNA sequence of shape (2048, 4) and the corresponding 4 bp resolution ATAC-seq of shape (512). To reduce the ATAC-seq resolution from 1 bp to 4 bp in the dataset, the values were binned using the maximum value.

Each model was trained on whole-genome data, excluding samples at genomic locations with unreliable accessibility measurements to reduce noise and biases. Specifically, we excluded samples overlapping with blacklisted regions (https://storage.googleapis.com/basenjibarnyard2/hg38.blacklist.rep.bed) or where less than or equal to 35% of the 2048 bp input overlapped with unmappable regions (https://storage.googleapis.com/basenjibarnyard2/umapk36t10l32hg38.bed). The annotations for blacklisted and unmappable regions of the hg38 genome were sourced from Kelley *et al* [2]. Reverse-complemented DNA sequences were augmented to the training data to enhance model robustness.

During the evaluation, we imposed a tighter constraint on the mappability and only evaluated samples that did not overlap with any unmappable region. Additionally, we used only the predictions corresponding to the central 1024bp genomic section. This ensured that there was at least 512 bp of genomic context present on either side of any region that we were trying to predict.

In addition to evaluating performance on the whole genome data, we also composed a dataset of peaks by taking samples centered on the ATAC-seq peaks to compare how models perform on high signal regions.

### Training details

We did a 5-fold cross validation where the genome was randomly split into 5 sets of chromosome – [1, 11, 13, 20], [2, 10, 14, 19, 21], [3, 12, 16, 17, 22], [6, 7, 9, 15], [4, 5, 8, 18]. For each fold, a model was trained for a maximum of 70 epochs with early stopping, with a patience of 5 epochs based on Pearson’s correlation on validation data.

Poisson negative log-likelihood loss was used to train all models with the AdamW [40] optimizer. We began by linearly increasing the learning rate during the first epoch until we reached a maximum of 1 *×* 10^*−*3^. We then used exponential decay coupled with cosine annealing [41], restarting at the beginning of every epoch.

### Model architectures

All models processed one-hot encoded DNA sequences of 2048 bp and predicted ATAC-seq signals corresponding to the central 1024 bp at 4 bp resolution. Each model is illustrated in Supplementary Fig. S1. The runtime and memory requirements of each model are described in Fig. 2d and Supplementary Fig. S2.

### CNN

The CNN, inspired by DeepSEA [1], was adapted for 2048 bp inputs and outputs. It comprised three convolutional layers with 512, 640, and 768 filters (kernel size 8), each followed by max pooling (window 4) and dropout (rate 0.2) to reduce dimensionality and prevent overfitting. The flattened output fed into two dense layers with 1024 and 512 ReLU-activated units. This architecture excels at detecting local motif patterns, such as transcription factor binding sites, indicative of chromatin accessibility.

### LSTM

The recurrent model (LSTM), based on DanQ [14], processed inputs through a convolutional stem with 512 filters (kernel size 26), followed by max pooling (window 4) and dropout (rate 0.2). A bidirectional LSTM layer [42] with 512 units captured longrange dependencies in the sequence. Two dense layers (1024 and 512 ReLU-activated units) produced the final output. The LSTM’s memory enables modeling sequential relationships across the DNA.

### dCNN

The dilated CNN (dCNN), inspired by Basenji [2] and adapted by ChromBPNet [4] for 2048 bp inputs, used 11 residual blocks. Each block contained two convolutional layers, the first with a dilation rate increasing by a factor of 1.5 across blocks, followed by GELU activation [43]. A dense layer reduced the channel dimension to 1, and a softplus activation ensured positive outputs. This design captures long-range genomic interactions.

To enhance feature extraction, we replaced traditional convolutional blocks with ConvNeXt V2 blocks [23], which offer improved efficiency and expressiveness via depthwise convolutions and layer normalization.

### ConvNeXtCNN

This model used a ConvNeXt stem with 128 filters (kernel size 15), followed by three ConvNeXt blocks with 256, 512, and 512 filters (kernel size 15), each applying depthwise convolutions [44]. Max pooling (window 4) and dropout (rate 0.2) followed each block. Two dense layers (512 and 32 ReLU-activated units) processed the feature matrix, with a softplus activation ensuring positive outputs.

### ConvNeXtLSTM

The ConvNeXtLSTM employed a ConvNeXt stem with 256 filters (kernel size 26), followed by a bidirectional LSTM layer [42] with 256 units to model distal genomic interactions. The final LSTM state fed into two dense layers (256 and 512 ReLU-activated units), with a softplus activation for the output.

### ConvNeXtDCNN

Similar to dCNN, the ConvNeXtDCNN used 11 dilated residual blocks but replaced the initial convolutional layer with a ConvNeXt stem (256 filters, kernel size 15). The remaining architecture, including dilation rates and softplus output, mirrored dCNN.

### ConvNeXtTransformer

We introduced the first transformer-based model for ATAC-seq prediction. Five ConvNeXt blocks (256 filters, kernel size 15) and pooling layers tokenized the input into 512 sequences at 4 bp resolution. This sequence of 512 tokens was then passed through four transformer blocks. Mirroring the behavior seen in current large-language models [45, 46], we observed better performance when using rotary positional encodings (RoPE) [28] compared to conventional sinusoidal encodings proposed in Vaswani *et al* [47]. A dense layer aggregated the (512, C) output across channels, followed by a softplus activation.

### ConvNeXt-xLSTM

Lastly, we built an extension of the above methods by using xLSTM in the core of the model. xLSTM is a newly built model that extends LSTM by retaining memory across different states, thereby capturing both local and global contexts [29]. The input DNA sequence was passed through 5 ConvNeXt blocks followed by an xLSTM block consisting of three mLSTM and one sLSTM blocks with a kernel size of 4 and 4 heads (Supplementary Fig. S3). The output was then passed through a dense layer with 128 hidden units and softplus activation function.

### Hyperparameter tuning

To optimize model performance for ATAC-seq signal prediction, we tuned hyperparameters for each method by maximizing Pearson’s correlation on the validation set of fold 0, comprising held-out chromosomes. For all models, we tested batch sizes of 64, 128, and 256, initial learning rates of 0.005, 0.001, 0.0005, and 0.0001, and a cosine annealing scheduler with warmup ratios of 0.1, 0.25, or 0.5 of total epochs [41].

For CNN-based methods (CNN and ConvNeXtCNN), we tuned the number of convolutional layers in the core block (2, 3, or 4), the kernel size in the core block (3, 7, or 15), and the number of dense layers in the output block. All combinations of these parameters were tested. Increasing the dense layers from 2 to 3 did not improve performance, so we retained 2 layers.

For LSTM-based methods (LSTM and ConvNeXtLSTM), we tuned the kernel size of the initial convolutional stem (3, 7, 15, or 26, inspired by DanQ [14]), the number of bidirectional LSTM layers (1 or 2), the number of dense layers (2 or 3), and their hidden sizes (256, 512, or 1024).

For dilated-CNN-based methods (dCNN and ConvNeXtDCNN), we explored architectures with 7 to 15 convolutional residual blocks, inspired by Enformer [3] and ChromBPNet [4]. We varied the dilation rate (1 to 2, in steps of 0.1) and tested three convolutional block configurations within each residual block. These configurations used kernel counts of 128 to 512 (step size 64), 64 to 256 (step size 64), or 256 to 4096 (step size 256), with corresponding kernel sizes of 5 to 20, 2 to 5, or 1 to 4, respectively. For the ConvNeXtTransformer, we tuned the ConvNeXt stem with 1 to 4 layers, kernel sizes of 7, 11, 15, or 26, and filter counts of 64, 128, or 256. The core ConvNeXt blocks were tested with 2 to 6 layers, kernel sizes of 3 to 11, and inverted bottleneck scales of 0.5, 1, 2, or 4, which adjust channel expansion in ConvNeXt’s depthwise convolutions [23]. The transformer core was optimized with 2 to 6 layers and feedforward dimension scales of 0.5, 1, 2, or 4.

### Adding mappability information to models

Experimental ATAC-seq signals are influenced by the mappability of the region of interest. While deep learning models can implicitly learn mappability from DNA sequences, explicitly providing this information can enhance certain models. To achieve this, we predicted mappability values from the initial layers of each model alongside the 4 bp resolution ATAC-seq signal (Supplementary Fig. S6).

The mappability values were calculated as a binary vector based on overlap with Basenji’s unmappable regions [2]. Zeros were assigned to the genomic segments overlapping with unmappable regions, and ones were assigned otherwise.

Adding this mappability information stabilized CNN and LSTM models but had no significant effect on dCNN or ConvNeXt-based models (Supplementary Fig. S6). This suggests that architectures capturing long-range contextual information, such as ConvNeXt, may implicitly encode mappability due to their larger receptive fields.

### Peak calling

Predicted ATAC-seq signals from each model were saved as BigWig files using the pyBigWig python package [48]. These were then converted to bedGraphs using bigWigToBedGraph from UCSC Genome Browser utilities [49] and sorted by chromosome and start position. Finally, peaks were called from the sorted bedGraphs using MACS2 bdgpeakcall command (version 2.2.9.1) [50] with a minimum score threshold of 1 to filter low-confidence regions.

### Variation score

To calculate the robustness of each model to input shifts, we used the variation score as described in Toneyan *et al* [21]. For each 512 bp region in our test dataset, we constructed 17 distinct 2048 bp long input sequences such that the output corresponding to each input contains the predictions for the selected 512 bp region. Using these 17 predictions, the coefficient of variation was calculated for each 4 bp bin, *i*.*e*. for each 4 bp bin, we calculated the mean and standard deviation of the predictions. The coefficient of variation for a given bin was then calculated as the division of the standard deviation by the mean across the 17 predictions.

Finally, the variation score was obtained by taking the average coefficient of variation across all the bins. For the whole-genome evaluation, this was repeated for all the 512 bp regions of the test chromosomes whereas for peak-centered evaluation, only those 512 bp regions were selected that contained an ATAC-seq peak.

### SNV filtering

We define a single-nucleotide variant (SNV) as either a somatic mutation or a singlenucleotide polymorphism (SNP). For the four primary tumor samples, we obtained somatic mutation calls from the study of pan-cancer analysis of whole genome (PCAWG) [51]. The SNP calls were obtained from GDC data portal [36].

For each patient, the SNVs were filtered to retain only those SNVs that significantly changed the chromatin accessibility profiles. First, somatic mutations that did not pass the quality control measures of PCAWG were removed. Similarly, we only selected those SNPs that had a heterozygous genotype and a high confidence score (uncertainty value *<* 0.1). The filtered files were then combined, and total read counts from the whole genome sequencing (WGS) and ATAC-seq experiments were calculated for the reference and variant alleles for each SNV. Using these values, the variant allele frequency (VAF) for each SNV was calculated, giving us WGS VAF and ATAC VAF. As we estimated the expected change in ATAC-seq signal for each SNV using the ATAC-seq read counts, we expected the SNVs to be unbiased towards either allele. Therefore, we selected SNVs with WGS VAF in the range of [0.4, 0.6]. Finally, to obtain SNVs that have a significant change in ATAC-seq, we calculated the significance of change using the Fisher exact test and the odds ratio. Each of these quantities was calculated using the reference and alternate allele read counts in WGS and ATAC-seq experiments. SNVs that had a p-value *<* 0.01 and odds-ratio *∈* [0.67, 1.33] were considered to have a significant change in ATAC-seq signal and therefore, were selected for SNV analysis.

This filtering led to a total SNV count of 1,025, with 80 somatic mutations and 945 SNPs across the four cancer patients.

## Declarations

### Funding

This project is partially funded by the Swiss Data Science Center (SDSC) collaborative projects grant (C22-09); Swiss Government Excellence Scholarship (ESKAS-Nr: 2021.0468) to A.G.

### Data availability

The input ATAC-seq bigwig or bam files were obtained from the ENCODE data portal [33] or the GDC data portal [36]. Each patient’s SNPs were also obtained from the GDC data portal [36], whereas information on somatic mutations was downloaded from the ICGC data portal [37]. All accession codes are provided in Supplementary Table S1.

### Code availability

The code for the ASAP framework has been made available at https://github.com/BoevaLab/ASAP.

## Notes

### Competing Interest Statement

The authors have declared no competing interest.

### Summary of Updates

The text is updated to be clearer. The figures are updated. The codebase is released. Supplementary figures and tables are added.

