## Supplementary Figures for "Early feature extraction drives model performance in high-resolution chromatin accessibility prediction: A systematic evaluation of deep learning architectures"

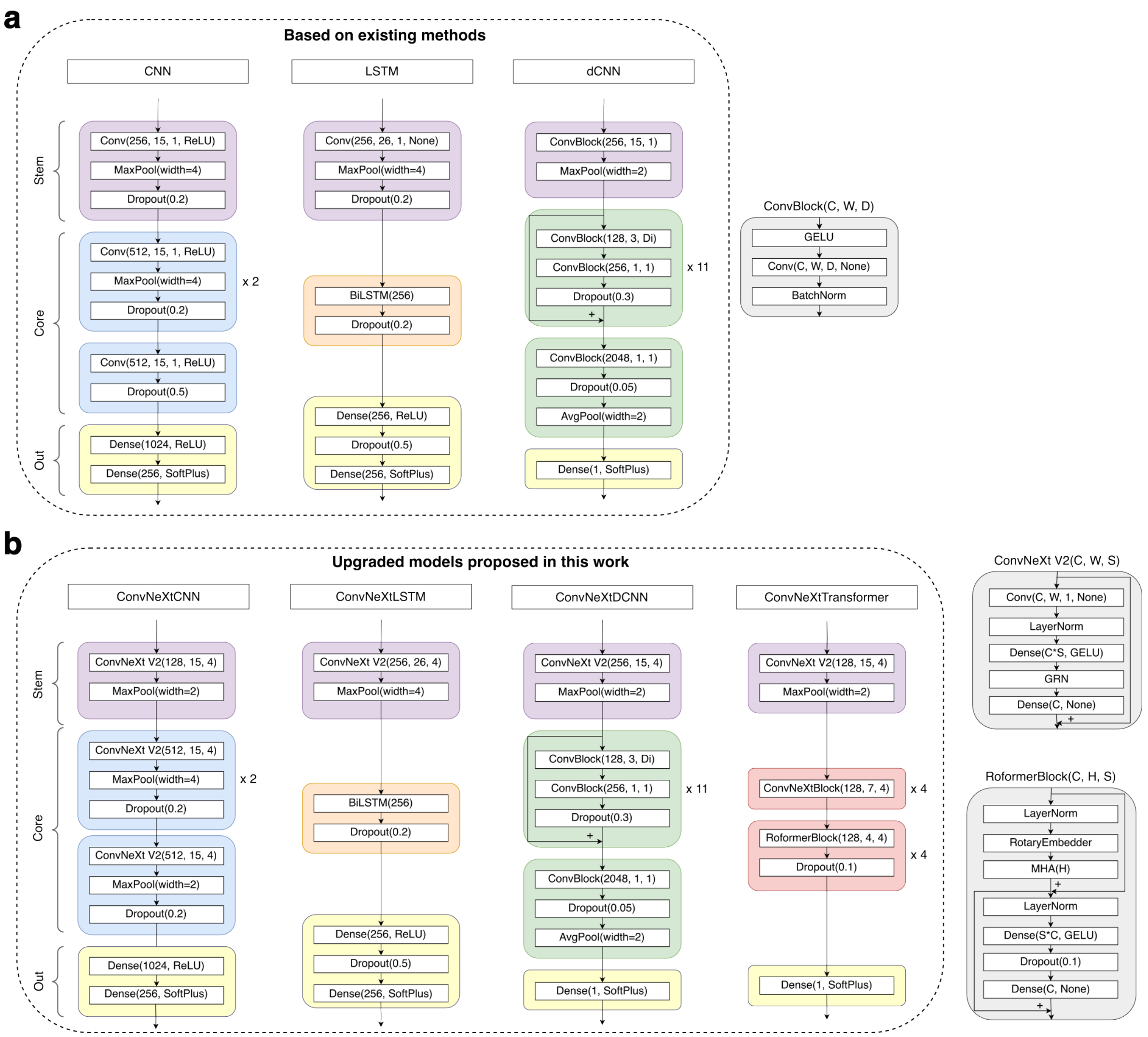


Figure 1: **Detailed model architectures. a,** The CNN, LSTM, and dilated CNN (dCNN) architectures based on existing works. **b,** The new models proposed in this work, including a transformer-based architecture. These models use ConvNeXt V2 block as feature extractors.

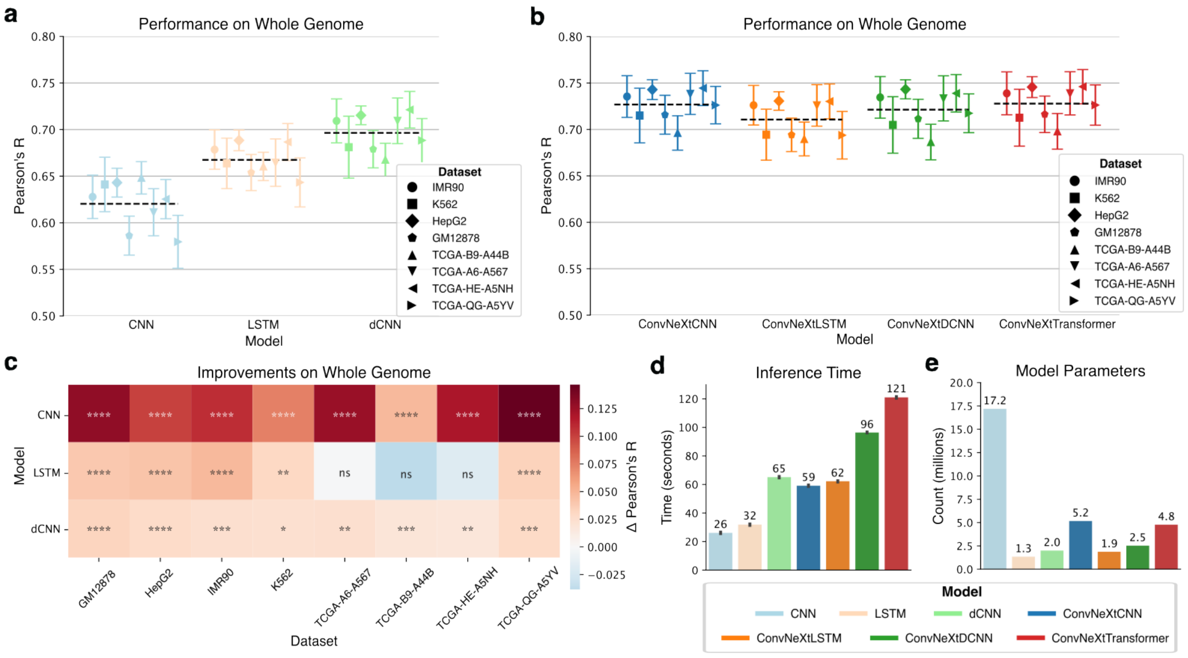


Figure 2: **Model comparison on whole genome.** **a,** Pearson's correlation between true and predicted ATAC-seq signals across whole genome of the test chromosomes across eight distinct datasets for state-of-the-art models. **b,** Performance comparison among the four models proposed in this work (ConvNeXtCNNs, ConvNeXtLSTMs, ConvNeXtDCNNs, and ConvNeXtTransformers) for the ATAC-seq peak regions stratified by cell lines and cancer patients. The black dashed line shows the average performance of a model across all datasets and chromosomes. **c,** Improvements of the new ConvNeXt-based methods proposed in this work as compared to existing methods. The significance is calculated with a two-sided Mann-Whitney U test on Pearson's R calculated for each test chromosome. ****: $P \leq 0.0001$, ***: $P \leq 0.001$, **: $P \leq 0.01$, *: $P \leq 0.05$, ns: $P > 0.05$. The $\Delta$ Pearson's R is calculated as the difference between mean Pearson's R across all chromosomes for a ConvNeXt-based method and the corresponding existing method. **d,** The total inference time in seconds for predicting whole genome region of chromosome 17 on a single RTX2080ti GPU for each model. **e,** Each model's trainable parameter count (in millions).

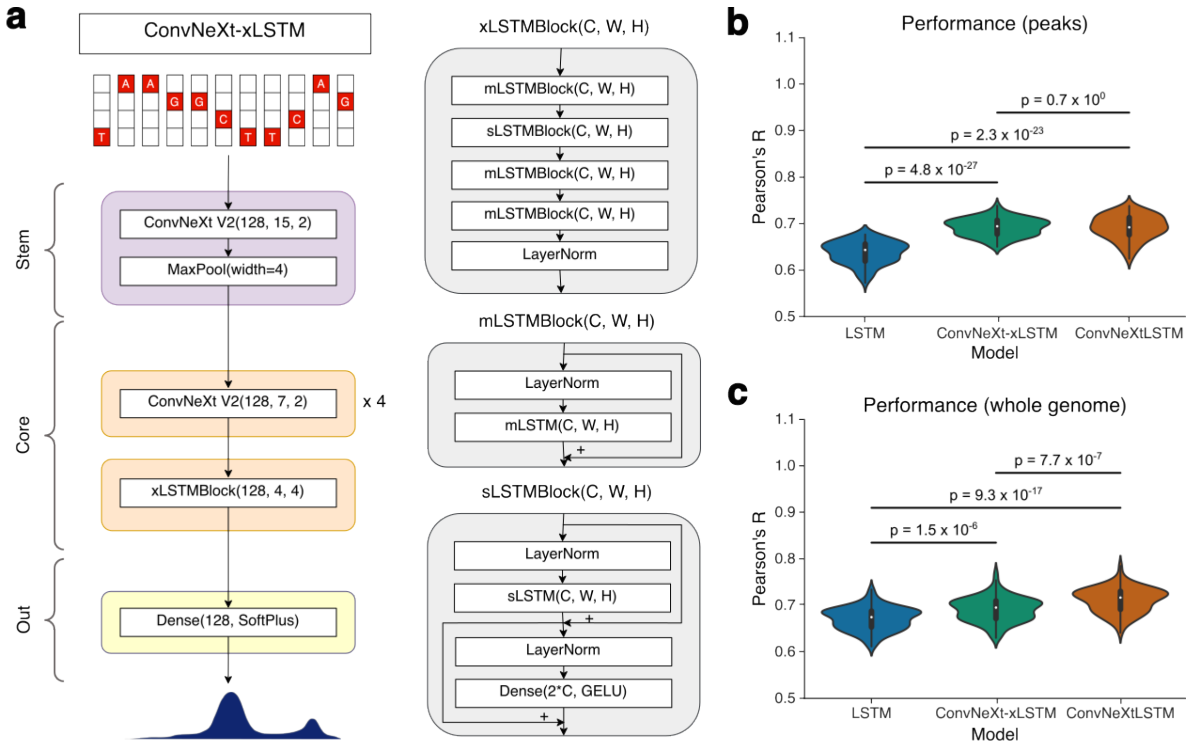


Figure 3: **Performance of xLSTM-based ATAC-seq prediction model. a,** We introduced ConvNeXt-xLSTM, which uses the newly designed xLSTM as the model core, for our task of ATAC-seq prediction. **b-c,** Comparing ConvNeXt-xLSTM against LSTM and ConvNeXt-LSTM based on Pearson's R on peak regions as well as whole genome of test chromosomes. The significance is calculated with a two-sided Mann-Whitney U test.

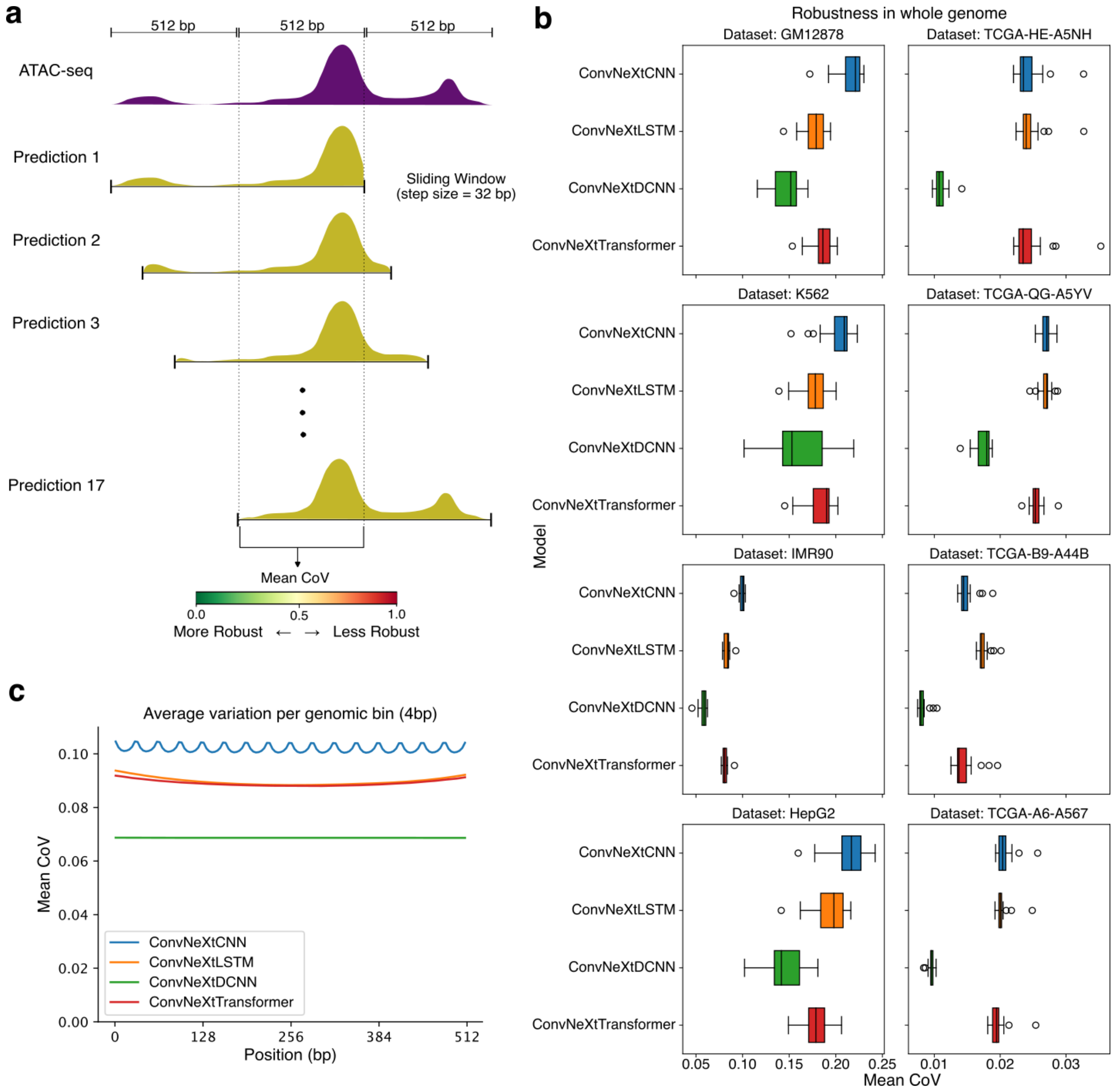


Figure 4: **Robustness test on whole genome.** **a,** The robustness test evaluates each method's ability to predict the same accessibility signal despite small shifts in the DNA sequence input. For a given model, the input is shifted by a few base pairs and the common predicted outputs are compared for variation. $N$ such predictions are taken into account by taking a sliding input window with a fixed step size. For our experiment, we choose $N = 17$ leading to a step size of 32bp. Variation is calculated as mean coefficient of variation (CoV) across all non-overlapping 4bp bins corresponding to the common 512bp genomic region. Lower CoV suggest high robustness to input shifts. **b,** The mean CoV measure for whole genome of test chromosomes across the eight datasets used in this study. **c,** Position-stratified mean CoV computed for each method across all the datasets.

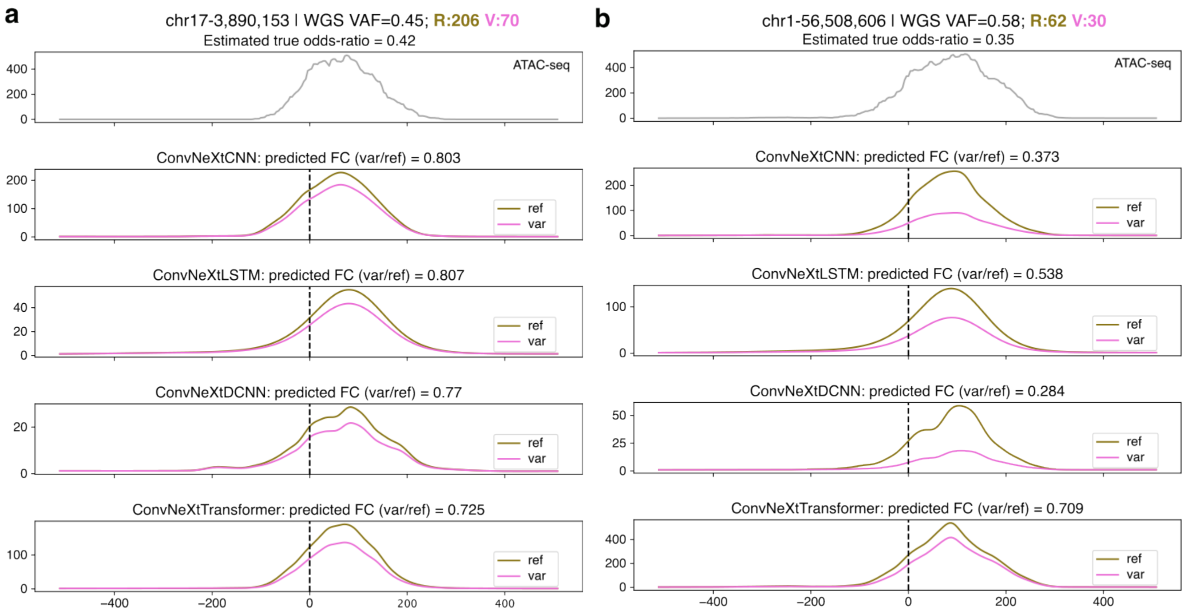


Figure 5: **Examples of allele-specific ATAC-seq prediction. a-b,** ATAC-seq prediction by our proposed methods for reference allele and genomic variant in chromosomes 17 and 1 respectively. WGS VAF: Whole genome sequencing variant allele frequency; R, V: total ATAC-seq reference reads and variant reads respectively.

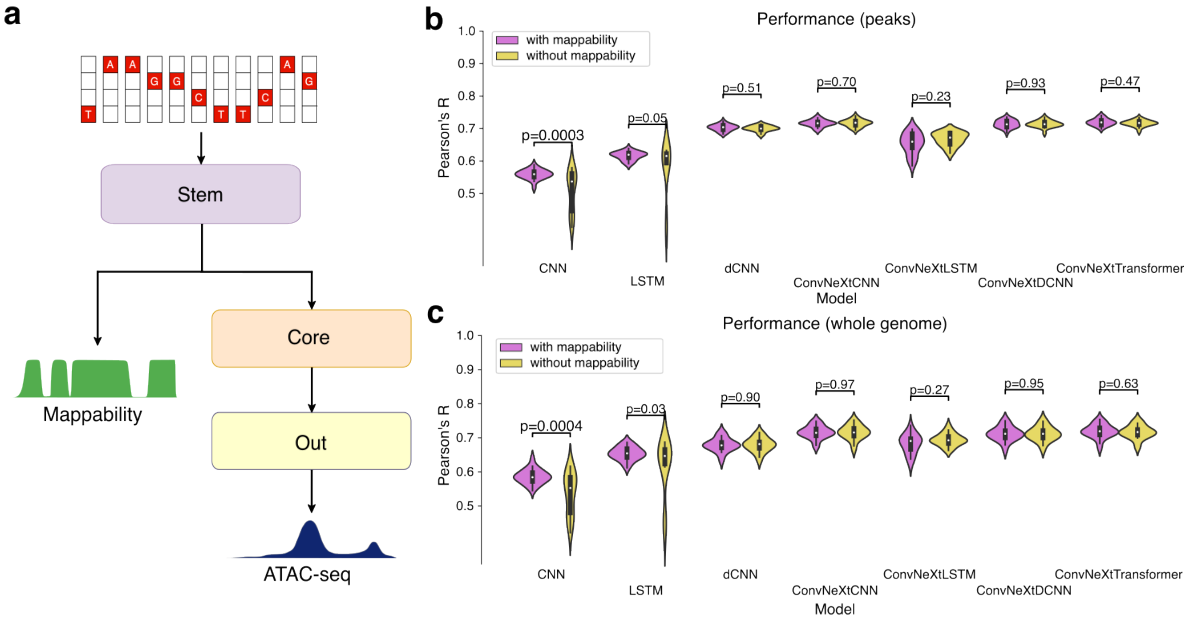


Figure 6: **Effect of including mappability information to the models. a,** Mappability values are additionally predicted by each model using the outputs of their stem block. **b-c,** Improvements in model performance are compared in peak regions and whole genome of test chromosomes caused by the addition of mappability information. The significance is calculated with a two-sided Mann-Whitney U test.
